# A new method for rapid genome classification, clustering, visualization, and novel taxa discovery from metagenome

**DOI:** 10.1101/812917

**Authors:** Zhong Wang, Harrison Ho, Rob Egan, Shijie Yao, Dongwan Kang, Jeff Froula, Volkan Sevim, Frederik Schulz, Jackie E. Shay, Derek Macklin, Kayla McCue, Rachel Orsini, Daniel J. Barich, Christopher J. Sedlacek, Wei Li, Rachael M. Morgan-Kiss, Tanja Woyke, Joan L. Slonczewski

## Abstract

Current supervised phylogeny-based methods fall short on recognizing species assembled from metagenomic datasets from under-investigated habitats, as they are often incomplete or lack closely known relatives. Here, we report an efficient software suite, “Genome Constellation”, that estimates similarities between genomes based on their k-mer matches, and subsequently uses these similarities for classification, clustering, and visualization. The clusters of reference genomes formed by Genome Constellation closely resemble known phylogenetic relationships while simultaneously revealing unexpected connections. In a dataset containing 1,693 draft genomes assembled from the Antarctic lake communities where only 40% could be placed in a phylogenetic tree, Genome Constellation improves taxa assignment to 61%. It revealed six clusters derived from new bacterial phyla and 63 new giant viruses, 3 of which missed by the traditional marker-based approach. In summary, we demonstrate that Genome Constellation can tackle the computational and algorithmic challenges in large-scale taxonomy analyses in metagenomics.

## INTRODUCTION

The McMurdo Dry Valleys, Antarctica, is one of Earth’s coldest dry deserts. Despite these year-round harsh conditions, this unique landscape harbors numerous perennially ice-covered, permanently meromictic lakes with unique geochemistry and microbial communities dominated by the microbial loop (Spigel and Priscu, 1998; Priscu et al., 1999; Green and Lyons, 2009). Several of the lakes have been intensively studied for more than 3 decades as part of the McMurdo Long Term Ecological Research Program (`mcmlter.org`). Along the valley, 80 km across the Ross Sea from McMurdo Station, Lake Fryxell is located at the lower end of Taylor Valley and situated between two glaciers, the Canada and the Commonwealth. Lake Fryxell is relatively shallow (20 m) and fed by 13 ephemeral streams for 4-12 weeks during the summer months. Another lake in the Taylor Valley is Lake Bonney. Lake Bonney is located at the western end of Taylor Valley and divided into two lobes (west and east) separated by a narrow sill which allows mixing between the lobes for a few weeks during the austral summer. The west lobe of Lake Bonney is connected to Taylor Glacier which harbors a unique geochemical feature, Blood Falls (Mikucki et al., 2009). Lake Bonney is connected to significantly fewer streams and is significantly deeper (40 m) than Lake Fryxell. The water columns are dominated by the microbial loop, with few to no metazoans (Priscu et al., 1999). The water columns of these lakes exhibit permanent chemical and physical stratification, harboring unique bacterial and eukaryal communities adapted to extreme environmental conditions including oligotrophy, low temperature, and hypersalinity (Roberts et al., 2004a; Li et al., 2016b; Kwon et al., 2017). In addition, thick benthic cyanobacterial mats carpet the lake floors to the base of the photic zone, often forming complex pinnacle-shaped structures reaching several centimeters tall (Jungblut et al., 2016; Hawes et al., 2013). As the lakes and their ecosystems have been isolated for hundreds of years, they provide an excellent opportunity to study the phylogenetic diversity change of the microbial species in response to climate change (Hall et al., 2017).

Microbial species classification has evolved from phenotypic description to genetic characterization, from pairwise genomic DNA–DNA hybridization to 16S ribosomal RNA (rRNA) homology (for a comprehensive review please see (Achtman and Wagner, 2008)). With the advent of next-generation sequencing and its application to metagenomics, we now have an efficient toolset to rapidly identify hundreds of thousands of microbial species from a single sample. Many of these microbial species are novel and have not been previously identified (Pasolli et al., 2019). More robust classification methods have been developed to consider multiple phylogenetic markers (Wu and Eisen, 2008) (Waterhouse et al., 2012), or even the entire genome (Rannala and Yang, 2008). Compared to other alternatives, whole genome-based classification methods carry the least biases and has the potential to discover novel species that share little homology to existing ones in current databases, such as in the case of giant viruses (La Scola et al., 2003).

Traditional whole genome-based phylogenetic methods rely on sequence alignments. They first identify homologous regions between two genomes, at either the DNA or protein level, and then use these alignments to build phylogenetic trees to infer relationships (Konstantinidis and Tiedje, 2005)(Parks et al., 2018). This approach has several limitations, including slow computing, low scalability, and low sensitivity to detecting remotely related species (reviewed in (Zielezinski et al., 2017)). To overcome these limitations, several alignment-free methods have been developed that are both computationally efficient and resistant to noise (Zielezinski et al., 2017; Ren et al., 2018; Jain et al., 2018). Based on their methodologies, there are three main categories of alignment-free genome similarity comparisons (reviewed in (Zielezinski et al., 2017)). Briefly, k-mer/word-frequency-based methods first compute a vector of frequencies for all or a subset of short k-mers (words) for each genome, and similarity is calculated using specific similarity functions. Substring-based methods estimate genome similarity using shared substrings, while information theory-based methods use sequence complexity or entropy to estimate their similarities. Recently, machine learning-based methods have also been developed that either use species-specific k-mers (such as Kraken (Wood and Salzberg, 2014) and Taxonomer (Flygare et al., 2016)) or secondary features derived from primary sequences (Li et al., 2017) for species classification.

As the computing requirement for pairwise genome comparisons grows quadratically with the increasing number of genomes, alignment-free methods that rely on a full set of k-mers become time- and cost-prohibitive with hundreds of thousands of genomes. One strategy to overcome this limitation is to use a small set of k-mers. For example, Mash only uses a small set of random k-mers (with smallest hash values, or MinHash) to approximate the similarity between sequences with only minor accuracy loss (Ondov et al., 2016). Minimizers, on the other hand, chooses the smallest k-mers in alphabetical order within a window (Roberts et al., 2004b). Most recently, advanced indexing methods that enable ultra-fast searches at the read level have also been developed (Bradley et al., 2019). These heuristic algorithms have demonstrated dramatic speed and space advantages; however, their accuracy suffers when dealing with poor genome assemblies from metagenomes (Olm et al., 2017). In addition, none of the current solutions have a built-in function to visualize the relationships between genomes, which could be very useful for detecting unusual patterns. Visualization of genome similarities so far has been largely restricted to tree-based methods, and these methods are not suitable for large numbers of genomes, nor do they offer interactive exploration capabilities.

In this study we developed Genome Constellation, a set of tools for the analysis of draft genomes assembled from metagenomes with the following functionalities: 1) A Bit vector implementation for fast genome similarity comparison; 2) genome similarity-based clustering; 3) k-nearest-neighbor (KNN)-based taxonomy classification; and 4) A web-based interactive visualization tool. We demonstrate these functionalities using two reference genome datasets and a set of metagenome-assembled genomes (MAGs) from the Antarctic lake communities. The software is available under BSD license at https://bitbucket.org/berkeleylab/jgi-genomeconstellation/.

## METHODS AND MATERIALS

### Using bit vectors as genome fingerprints, or Spectrum

To transform a genome sequence into a bit vector, we hash the k-mers of a genome into a vector of fixed length *n* (the spectrum vector) as follows. First, each of the k-mers is hashed to a 128-bit k-mer vector using MurmurHash 3. We choose a fixed length *k*=23 as a trade-off between the ability to differentiate strains of the same species and sufficient sensitivity to detect genus level similarity. A bit-mask can be used to select a deterministic subset of k-mers, each masked bit cutting the fraction of selected k-mers in half. Second, the *p*-th bit of the *n*-bit spectrum vector is set to “on,” where *p* is the integer represented by the second half (64 bits) of the k-mer vector modulo *n*. The saturation of the bits set “on” in the spectrum can be controlled by varying the length of the spectrum and the size of bit-mask. The larger the length of the spectrum, the more bits are masked, the lower the saturation rate a spectrum has and vice versa. For example, a 10Mb genome has 10 million k-mers, if we sample 1 out of every 1000 k-mers (at most 10,000 unique k-mers) and the spectrum vector is 1 million in length, the resulting spectrum vector will have less than 1% bits are on, or 1% saturated. It is crucial to keep the saturation rate low to avoid hash collisions, i.e., different k-mers map to the same bit on the spectrum vector. These steps are illustrated in Figure 1A.

**Figure 1.**
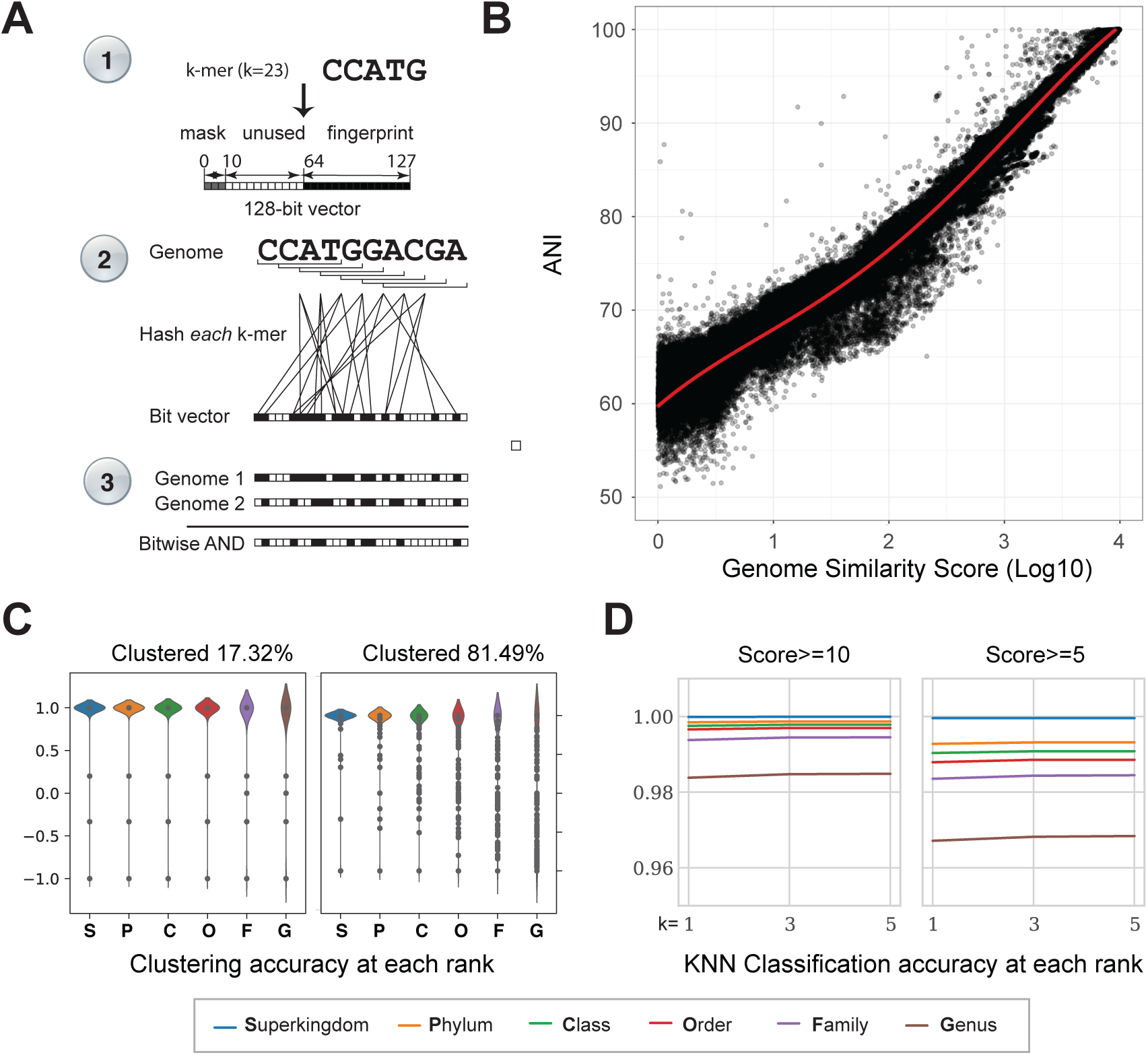
Using bit vectors to estimate sequence similarity. **(A)** We use three steps to compute the similarity between any pairs of genomes. First, we hash k-mers (default=23) to a 128-bit vector and use the first half as a mask for selecting a random set of k-mers (default=10 bits, or 1 in 1024), and the second half to determine the “on” bits in the genome spectrum vector in the second step. In the third step, the genome spectrums can be bitwise compared by the shared “on” bits, a very efficient ADD operation. See methods for details. **(B)** A scatter plot to compare the Average Nucleotide Identity (ANI) and Genome Constellation (GC) Score between all pairs of the NCBI 7k reference genomes. The x-axis is GC score at the log10 scale, and the y-axis is ANI (%), and a regression line is shown in red. **(C)** Clustering NCBI 7k reference genomes based on GSSs. (Left) At a stringent score cutoff (eps=20.0), 17.32% of the genomes were clustered. The clustering accuracy distribution is shown as violin plots at **S**uperkingdom, **P**hylum, **C**lass, **O**rder, **F**amily and **G**enus level, respectively. Except for a few outliers, clustering accuracy is almost perfect at all levels. (Right) At a less stringent score cutoff (eps=60.0), the clustering distribution was shown. At this cutoff, the majority of the genomes (81.49%) were clustered. Clustering accuracy is very good at Superkingdom, Phylum and Class levels. **(D)** *k*-nearest-neighbor classification performance with Genome Similarity Score (GSS) on 60k IMG known genomes. Leave-one-out classification experiments were carried out at a different number of neighbors (*k*, x-axes), and the accuracy is shown on y-axes, at two different GSS cutoffs (columns). Accuracy was measured at six different taxonomy ranks (superkingdom, phylum, class, order, family, and genus).

### Genome Similarity Scores (GSS)

A comprehensive list of existing methods of comparing the similarity between two bit vectors can be found in (Choi et al., 2010). Using the same notation, here *a* is the number of bits where the vectors are both 1s (or presence of the same k-mers in both genomes), meaning ‘positive matches,’ *b* is the number of bits where only the second vector are 1s (k-mers unique to genome 2), *c* is the number of bits where only the first vector are 1s (k-mers unique to genome 1), and *d* is the number of bits where both vectors are 0s (or absence of these k-mers in both genomes), meaning ‘negative matches.’ *n* is the sum of the four possibilities (*n* = *a* + *b* + *c* + *d*). Here we adopted a modified version of Forbes-II similarity, defined in Equation 1:

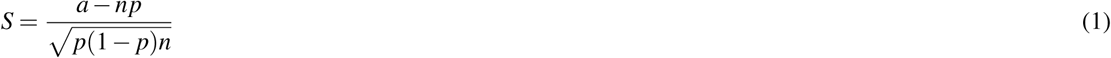

where *p* measures likelihood of k-mer matches between two unrelated sequences, and it is defined in Equation 2:

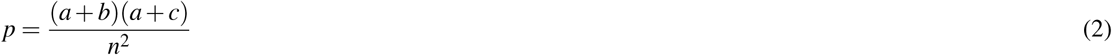

### Datasets used in benchmarking GSS

#### NCBI 7k Reference genome dataset

NCBI Reference genomes were downloaded from the NCBI FTP archive (as of June 2017). These include 29,195 Eukaryota, Bacteria, and Archaea genomes, 7,254 Virus and 84 Giant Virus genomes. For species with multiple strains, only one representative was randomly chosen to reduce redundancy. Genomes with a size smaller than 200kb or larger than 200Mb were also excluded. A total of 7,261 genomes remained, including 222 viruses (presumably giant viruses), 631 Eukaryota, 6,100 Bacteria and 303 Archaea genomes. In addition, 4 synthetic genomes (3 JCVI Mycoplasma and 1 from Synthetic Yeast Genome Project) where included. The complete list of this set can be found in Supplemental_File_1. This dataset is referred as “NCBI 7k Reference” throughout the text.

#### IMG100k genome dataset

A set of the sequenced microbial genomes were collected from the Integrated Microbial Genomes and Microbiomes (IMG/M) database (Chen et al., 2018) as of 09/19/2018. The same size filtering criteria were applied (between 200kb and 200Mb) to obtain 98,048 genomes. After excluding 15,557 unclassified genomes, the remaining set includes 79,290 Bacteria, 2820 Archea, 143 Eukaryota, and 238 Viruses. A complete list of these genomes and their taxonomy annotations is available in Supplemental_File_2. This dataset is referred as “IMG100k” throughout the text.

#### Antarctic genome dataset

Microbial communities were sampled from two meromictic lakes of the Taylor Valley, Victoria Land, Antarctica. Water was sampled from the permanent chemoclines of Lake Fryxell (77.605°S, 163.163°E, 9 m depth) and Lake Bonney, east lobe (77.719°S, 162.283 °E, 15 m depth). Water samples were concentrated by filtration through 0.45 *μ*m filter. DNA was extracted from the filters using an MP FastDNA SPIN DNA kit (MP Biomedicals, CA) following the manufacturer’s instructions (Bielewicz et al., 2011). Microbial lift-off mat samples were obtained from the surface ice of Lake Fryxell. Mat DNA was extracted by the PowerBiofilm kit (MO BIO). Shotgun metagenomic library construction and sequencing were carried out at DOE Joint Genome Institute (JGI) using standard protocols on the Illumina HiSeq 2500 platform. The raw reads were quality filtered using JGI standard protocols and 1.2Tb total sequences were obtained. The sequences and sample information are available at the JGI GOLD database with the accession no Gs0118069 (https://gold.jgi.doe.gov/study?id=Gs0118069).

The reads were assembled using Hipmer (Georganas et al., 2015) with default parameters except that the minimum contig size was set to 2.5kb. Unassembled reads were assembled using RAY assembler (Boisvert et al., 2010) with k-mer=53, and assembly of any yet remaining unassembled reads using Megahit (Li et al., 2016a) for a final assembly with the following parameters: k-list=33,43,63,83,103 min-contig-len=500bp. Genome bins were formed by MetaBAT2 (Kang et al., 2019) on contigs larger than 2.5kb (MetaBAT’s default) from the final assembly experiment. These bins or metagenome-assembled genomes (MAGs) are referred as “Antarctic MAGs” throughout the text. The assembly, bins, and associated analysis results are available for download at http://share.jgi-ga.org/?prefix=genome-constellation/.

### A WebApp for microbial genome visualization

We developed a Web App for searching similar genomes using Python/Flask and hosted on Amazon Web Service (AWS), Elastic Container Service (ECS). The URL of this app is http://constellation-classifier.jgi-ga.org. With this app, a user may upload one or more genomes and search for similar genomes in the NCBI 7k or IMG100k reference sets. Most searches only take a few seconds. The results from the NCBI 7k Reference can be directly sent to visualization (below).

We implemented a force-directed graph visualization (Frick et al., 1994) for Genome Constellation using a publicly available JavaScript library called vivagraph https://github.com/anvaka/VivaGraphJS. The web GUI uses vivagraph’s WebGL rendering and an initial antigravity force constant of −1.2. Tools to adjust the distance scaling factor and the antigravity force constant are provided to interactively optimize data distribution. An adjustable oval boundary is applied to prevent singletons from moving out of range. To improve performance, the genome data are divided into separate data files for node and link definitions, and the large link file can be further divided into smaller files. The data files are loaded concurrently using web workers to improve the loading time. Genome nodes are added to the graph first and simulation starts right after the nodes are all in the graph. Links are added in chunks while the simulation is running to achieve a pleasant visual effect.

### Classification based on GSS

The IMG100k reference genome set was used to evaluate classification performance as it is a comprehensive dataset. We selected 60,327 genomes that have known annotation at all ranks (superkingdom, phylum, class, order, family, genus, and species) for classification evaluation.

Taxonomy classification was carried out using the *k*-nearest-neighbors (KNN) algorithm, classification accuracy was calculated based on leave-one-out cross-validation. Briefly, in each iteration, the annotation of one of the genomes in the dataset was predicted based on the majority of its closest *k* neighbors at each rank, as measured by having the smallest GSS, and the prediction was compared to its true annotation. The percentage of correct predictions at each rank was computed after all the genomes had gone through one iteration.

### Co-clustering Antarctic genome bins with known reference

The GSS scores of the Antarctic genome bins and the NCBI 7k reference genomes were normalized and then converted to a sparse distance matrix. *de novo* clustering was performed using the DBSCAN function from scikit-learn v0.20.1. The sparse distance matrix was used as a precomputed matrix for clustering, with eps=60.0 and min samples=2.

Clustering performance was evaluated based on reference genomes. For each of the clusters, the mean score of all the pairs of genomes (1 when the pair comes from the same group at a particular rank, −1 otherwise) was computed, and the mean of the scores from all clusters was used as the overall clustering accuracy for each rank (superkingdom, phylum, class, order, family, and genus). For very large clusters, a random set of 100 genomes was chosen to estimate the mean cluster score (9,900 pairs).

### Antarctic MAG annotation

We used two different methods to quickly filter out known species. The first method is CheckM (Parks et al., 2015). Among the seven ranks (superkingdom, phylum, class, order, family, genus, and species) we only retained the highest taxonomy rank that CheckM was able to assign a genome bin to. The MAGs that CheckM was not able to assign any rank were labeled them as “na.”

We also developed a new method called LastTaxa for taxonomy classification of the Antarctic MAGs. For each MAG, we first predicted protein-coding genes using prodigal (http://prodigal.ornl.gov/) and then carried out a homology search against NCBI’s non-redundant database (NR) database using LAST (http://last.cbrc.jp/). Each of the proteins give a predicted taxonomy result based on its best hit (i.e. casts a vote), and the taxonomy predicted by the majority of the proteins (i.e. taxonomy receives the most votes) is then assigned to the genome bin. More details about LastTaxa are available at https://gitlab.com/jfroula/lasttaxa.

### Phylogenomics analysis

To place the Antarctic MAGs into an existing taxonomic framework of Bacteria and Archaea, MAGs were added to a representative set of microbial genomes available in IMG/M and the Genome Taxonomy Database (Parks et al., 2018). 56 universal single-copy marker proteins were used as phylogenetic markers (Rinke et al., 2013). MAGs with, MAGs were excluded from further analyses if they meet one of the following three conditions: 1) having less than 15 of the 56 unique markers, 2) having more than 5 different single-copy markers that have multiple copies, or 3) having one or more single-copy markers that have 5 or more copies. For every marker protein of the remaining genomes, an alignment was built with MAFFT (v7.294b) (Katoh and Standley, 2013) and subsequently trimmed with BMGE using BLOSUM30 (Criscuolo and Gribaldo, 2010). Single protein alignments were then concatenated and phylogenetic trees inferred with FastTree2 using the options: -spr 4 -mlacc 2 -slownni -lg (Price et al., 2010).

To infer the phylogenetic position of novel giant viruses, five core nucleocytoplasmic large DNA virus (NCLDV) proteins (Yutin et al., 2009) were selected: DNA polymerase elongation subunit family B (NCVOG0038), D5-like helicase-primase (NCVOG0023), packaging ATPase (NCVOG0249) and DNA or RNA helicases of superfamily II (NCVOG0076) and Poxvirus Late Transcription Factor VLTF3-like (NCVOG0262), and identified with hmmsearch (version 3.1b2, hmmer.org). Protein sequences were aligned using MAFFT (v7.294b)(Katoh and Standley, 2013). Highly gapped columns (less than 10% sequence information) were removed from the alignments with trimal v1.4 (Capella-Gutiérrez et al., 2009). Phylogenetic trees for each protein and for a concatenated alignment of all five proteins were constructed using IQ-tree (v1.6.10) with LG+F+R6 as suggested by model testing as the best-fit substitution model (Nguyen et al., 2015).

Phylogenetic trees of novel bacteria and virus clusters were generated using the Antarctic genome bins and reference genomes. This data was visualized using the Interactive Tree of Life (Letunic and Bork, 2019). Following the creation of these trees, Genome Constellation results from the Antarctic dataset were compared to the trees. This comparison assessed whether there were novel genomes found by Genome Constellation that were missed by the phylogenetic trees.

## RESULTS

### A genome spectrum for fast similarity estimation

To enable rapid estimation of pairwise similarity between thousands of genomes, we developed a new genome fingerprint, called spectrum, in the format of bit-vectors because bitwise computation is very efficient (Figure 1A and Methods). To increase sensitivity, we used all k-mers of a genome as its spectrum (full spectrum) to calculate the genome similarity scores (GSS) between genomes, and test how well they could be used to approximate evolutionary relationships between genomes.

Among existing methods to measure genome relatedness, Average Nucleotide Identity (ANI) has been used as a gold standard for closely related prokaryotic species (Konstantinidis and Tiedje, 2005; Varghese et al., 2015). To explore the relationship between ANI and GSS, we calculated both for a set of ∼7,000 genomes including prokaryotes, large viruses (>=200kb) and small eukaryotes from the NCBI RefSeq database (Methods). As shown in Figure 1B, Log-transformed GSS is closely correlated with ANI. In particular, the greater the ANI is, the stronger the linear relationship between them is. These results suggest that GSS accurately approximates ANI in measuring genome similarities.

Bitvector-based GSS calculation is very efficient. Comparing 100 genomes only takes one minute with 16-threads, roughly twice the speed of Mash (Ondov et al., 2016), 14x that of fastANI (Jain et al., 2018), and 66x that of the alignment-based MUMmer method under similar settings (Supplemental Script 1). One also has the option to use a random subset of the k-mers combined with a smaller spectrum size for GSS calculation (partial spectrum), which reduces sensitivity but gains much greater computational performance. With this option one can scale up to hundreds of thousands of genomes to rapidly detect highly similar genomes (e.g., ANI 85%, Supplemental Script 1). As expected, GSS does not change when it is applied to incomplete genomes, and only lateral transferred segments larger than 10kb can raise estimated ANI values consistently above 70% (Supplemental Script 1).

### Genome Similarity Scores can be used to rapidly cluster and classify species from different domains

The Genome Similarity Score (GSS), for the first time, puts all genomes into the same metric space, as it does not depend on any phylogenetic markers such as 16/18S ribosomal RNA genes or single-copy conserved genes. As such, it is conceivable that GSS could be used to effectively cluster and classify species from different domains.

GSS enables the unsupervised clustering of genomes into closely related groups. We first tested whether they can be used to cluster known genomes by running DBSCAN clustering algorithm on the set of 7k reference genomes. For each of the clusters formed, we designated each pair as 1 if it belongs to the same taxonomy group at a given level, and −1 otherwise. An average score between −1 and 1 was computed for each cluster to reflect the clustering accuracy and then they were averaged to represent the overall clustering accuracy. The distribution of clustering accuracy for each cluster at each taxonomy level is shown in Figure 1C. At stringent cutoffs where higher GSSs are used, fewer genomes are clustered and the cluster accuracy is almost perfect at all taxonomy levels.Conversely, cutoffs at lower GSSs result in more genomes being clustered, but the clustering accuracy decreases, especially at Order, Family and Genus levels.

As more and more genomes are sequenced and assigned taxonomy, it is possible that a non-parametric classifier, such as *k* -nearest-neighbor (KNN), could accurately predict the taxonomy of an unknown genome based on its similarity to known ones. We tested the above hypothesis on the IMG100K dataset (Methods). In each prediction, a genome’s taxonomy is assigned using the majority of its nearest neighbors (1, 3, 5, 7, 9). Stringency is controlled by a GSS cut-off parameter, which selects the similarity needed for a relationship to be considered. Additionally, GSSs are weighted using Shepard’s method (Shepard, 1968). The results are shown in Figure 1D.

The accuracy is the highest at the most stringent level (GSS cut_off=10) where 90.5% genomes have at least one neighbor. This results in an accuracy of more than 98% for each taxonomic rank. Using more neighbors decreases classification accuracy, but this can be corrected by using weighted GSSs to bias towards closer neighbors. Superkingdom-level classification accuracy is the highest, at almost 100%, while accuracy slightly decreases as the rank becomes more specific. This is likely to be explained by the fact that the number of genomes with the same group as the genome tested becomes fewer as rank increases. At a less stringent level, GSS cut _off=5, where 94.0% genomes have at least one neighbor, a similar trend is observed with a slightly lower classification accuracy, mostly at or above 96%. These results suggest GSSs can be used to accurately predict taxonomic information for genomes with many close known relatives.

It is worth noting that taxonomy classification based on GSSs is very fast, as 60,000 genomes can be predicted in several minutes on a common workstation.

### An interactive web tool designed to simultaneously visualize tens of thousands of genomes and explore their relatedness

Inspired by the force of gravity that positions the stars in the sky, we developed the Genome Constellation web portal, where GSS is used as “gravity” to lay out the genomes in a 2D space. Briefly, the layout is generated by a force-directed graph. In such a graph, genomes are nodes and edges are formed between genomes if their GSS exceeds a certain threshold. The relative positions of the genomes on the layout are determined by two basics forces: a default repulsion force (anti-gravity) and a gravitational force proportional to the GSS. The balance between these two forces eventually forms the final arrangement of all the genomes in the “universe,” with related genomes clustered into “constellations.”

The Genome Constellation web portal has the following designed features. First, the layout is completely data-driven, non-deterministic, i.e., different input data and different simulation experiments will give rise to different results. When the input data is large, different simulations tend to give very similar layouts. Second, it provides several parameters to produce user-defined optimal visualizations, including link weights, antigravity forces, minimum GSS cutoffs (MSC). Third, it provides quite a bit of interactivity: different ranks of taxonomy can be colored differently or toggled on/off, individual and multiple genome nodes can be searched and selected to examine their subnetworks. Finally, user data can be loaded into the application session through the import tool and visualized on the fly.

The built-in reference genome data set contains the NCBI 7k set with about 1 million links. The web application can handle up to 1 million genomes and 2 million links with reasonably smooth simulation and user interactivity on a typical client machine (tested on a Windows laptop with Intel i5-2400 CPU and 8GB memory). A demo video of Genome Constellation is available at http://bit.ly/JGIGenomeConstellation. Figure 2A provides two views of this Genome Constellation map, at phylum and class rank, respectively. Despite the graph being drawn without using any known phylogenetic information, the genome constellation largely resembles known phylogenetic relationships where related genomes are close to each other and form discrete groups.

**Figure 2.**
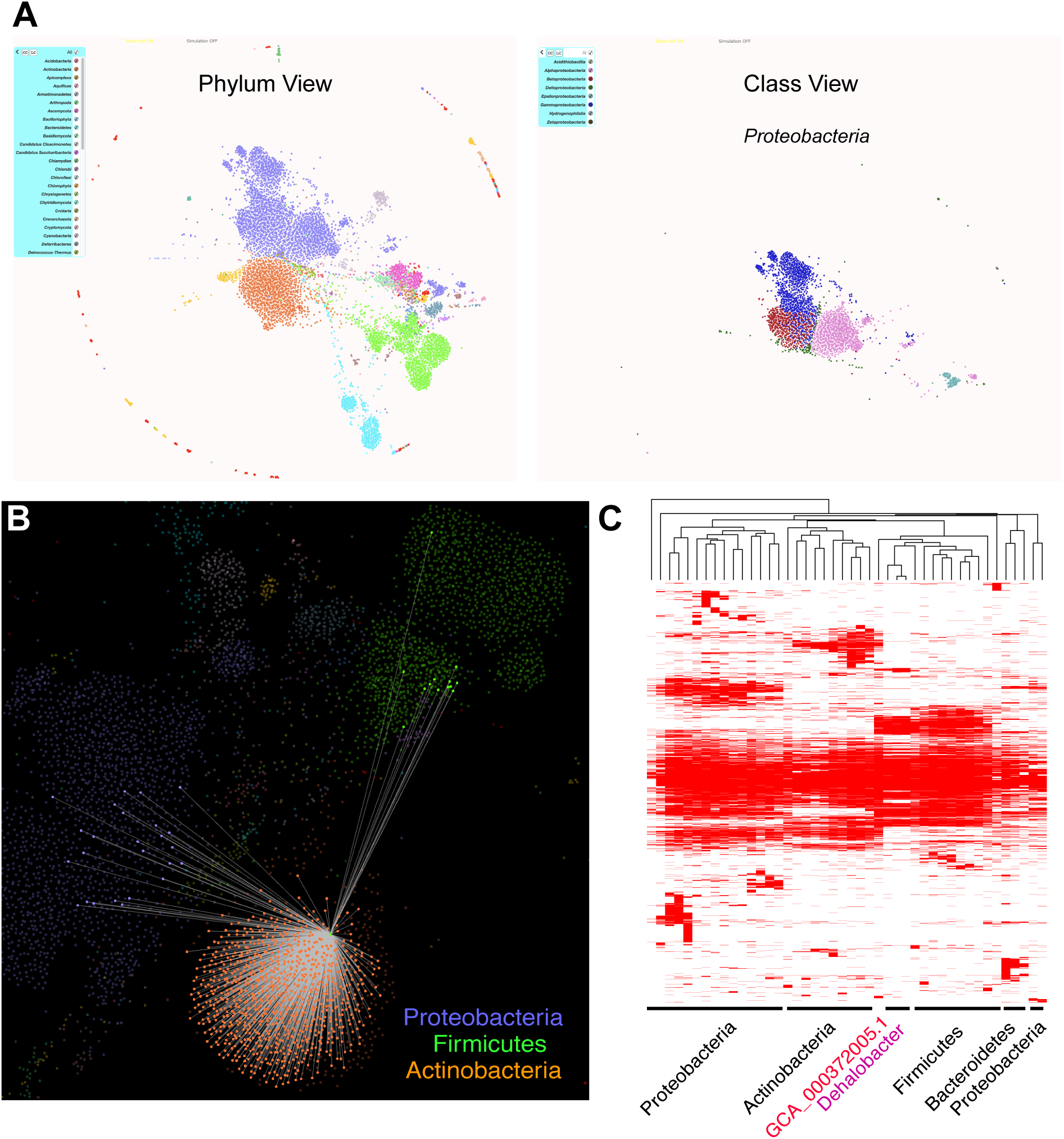
Genome Constellation Web App. **(A)**Two snapshots of Genome Constellation Map of 7,261 reference genomes from Antarctic mat and lake samples. (Left) A view at the phylum level, where each color represents a distict phylum. (Right) A view at the class level for Proteobacteria phylum. Each class is painted in a distinct color. Legends are shown on the left of each panel, and only a subset of the labels are shown due to the size limits. For the full legends please use the Web App. **(B)** Genome Constellation visualization reveals hidden relationships between known species. A known species, Dehalobacter sp. TeCB1 (GCA_001707455.1) that belongs to the Firmicutes phylum, but is placed within the Actinobacteria cluster. Its links to genomes from Actinobacteria (orange color), Firmicutes (green color) and Proteobacteria (blue color) are shown. **(C)** PFAM clustering analysis of Genome GCA_001707455.1 and its connected neighbors.

Genome Constellation can also uncover taxonomical errors and reveal hidden relationships among known genomes. It has been reported that current taxonomy assignment contains a substantial number of errors, either due to the manual process or lack of robust classification methods (Parks et al., 2018). For example, an Archaeal genome (GCA_000224475.1) annotated at the class level as a “halophilic archaeon” clusters with reference genomes in the species *Natronococcus occultus*, providing new and improved species identification for this genome.

Interestingly, a reference genome, GCA_001707455.1, annotated as one of the Firmicutes, is clustered closely with Actinobacteria, with some links to both Firmicutes and Proteobacteria (Figure 2B). The NCBI genome record suggests it is an unnamed species isolated from a population of Dehalobacter which are clearly distinct from the other species and was tentatively assigned as Dehalobacter (Tang et al., 2012; Deshpande et al., 2013). Moreover, protein family composition from GCA_001707455.1 resembles both Actinobacteria and Dehalobacter (Figure 2C), supporting this hypothesis that species represents a blend of a Firmicutes and an Actinobacteria. Its unique evolutionary history and metabolic capabilities would be very interesting to be further explored.

### Interrogate the Antarctica lake microbial communities with Genome Constellation

While the Taylor Valley lakes (Figure 3A) have been intensively studied for more than two decades as part of the McMurdo Long Term Ecological Research program (mcmlter.org), molecular genetic data on Dry Valley microbes remained limited until recently (Kwon et al., 2017; Taton et al., 2003; Vick-Majors et al., 2014; Li and Morgan-Kiss, 2019). In December 2014, the austral summer, we collected water samples from the chemoclines of Fryxell and Bonney (representing zones of high diversity and maximum productivity), and uplift mat samples emerging from the ice cover of Fryxell (Figure 3B). We carried out deep metagenome sequencing and assembled the 1.2Tb sequence data into 1,764 MAGs (483 and 1,281 from Bonney and Fryxell, respectively, see Methods). Among them, 71 MAGs were subsequently excluded from further analyses due to suspected high contamination rates.

**Figure 3.**
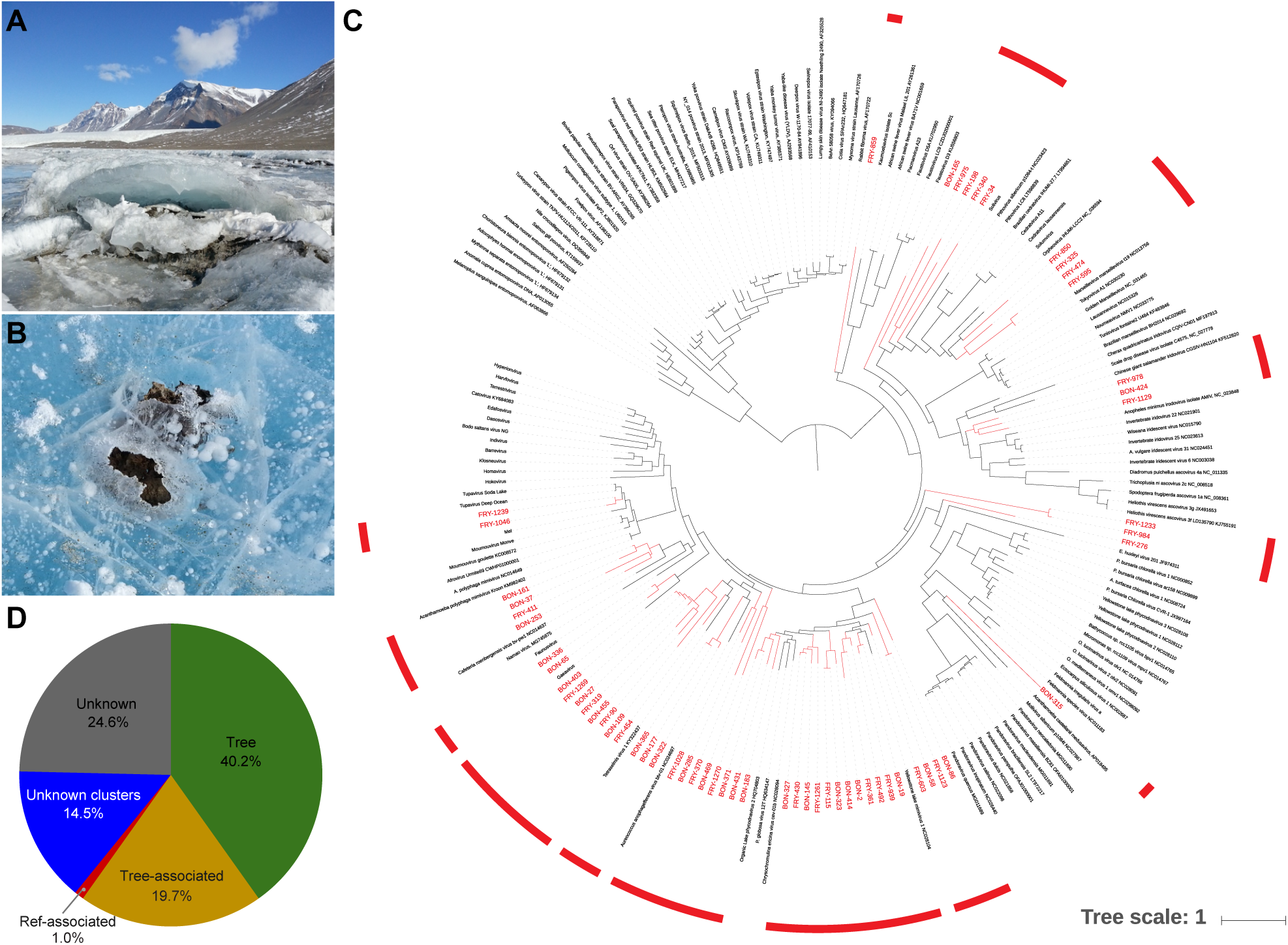
Clustering the Antarctic metagenome bins together with the 7k Ref genomes using GSS improves their taxonomy classification. **A**. Lake Fryxell with permanent ice cover, weathered by wind-driven sediment deposition and seasonal ablation of the ice cover; December 10, 2014. **B**. Uplift mat emerges from the ice; approximately 20 cm across. **C**. A phylogenetic tree constructed from 120 giant virus known references with the 60 Antarctic MAGs predicted by marker-based approach (red). **D**. Taxonomy classification coverage statistics for the phylogenetic tree-based (Tree), Clustered with tree-based (Tree-associated), Clustered with NCBI ref7k only (Ref-associated), Unknown clusters and Not clustered unknowns (Unknowns).

We first carried out phylogenomics analysis on the 1,693 MAGs, and were only able to place 680 (40.2%) on a Bacteria/Archaea or a Giant Virus phylogenetic tree (Methods, Figure 3C, and Supplemental Figure 1). This surprisingly low rate suggests these MAGs are either incomplete or derived from very novel taxa groups, or both. In addition, they could be Eukaryotic genomes that were not included in the tree-based analyses. The ability of Genome Constellation to cluster large numbers of genomes could be particularly useful for characterizing these MAGs in two ways: first, incomplete MAGs from known taxa groups could be recognized by their association with more complete members of the same cluster; Second, clustered MAGs from the same unknown taxa groups could increase the chance for detecting their remote homologs.

Expecting many novel taxa at the class level or higher from these isolated communities, we set a loose distance cutoff (eps=60.0 in DBSCAN) and clustered the Antarctic genome bins together with the 7k Reference genomes. At this cutoff, 1,172 total clusters with more than two genomes or MAGs were formed. At the phylum level, the clustering accuracy for reference genomes is 98.01+/-0.16%. 1,020 MAGs are clustered with or without known references. Among them, 235 MAGs (23.0%) are clustered with at least one known reference, including 136 that are missed by the phylogenomics analysis. 785 MAGs (77.0%) are not clustered with any of the known reference genomes in the NCBI 7k set, including 118 in completely unknown clusters (three or more). Overall, while tree-based analyses only covered 40.2% of the MAGs, clustering either with a MAG in the trees or with a known reference expanded the coverage by 19.7% and 1.0%, respectively. Among the remaining MAGs not covered, 14.5% form clusters while 24.5% are not covered by either the tree or the clustering experiments (Figure 3D). The expanded 20.7% taxonomy assignment of the MAGs also increase the mapped short reads from 1.04 billion to 1.30 billion (by 25.29%), which should improve downstream analyses based on species abundance.

We further characterized the 25 unknown clusters by first predicting encoded proteins of each MAG and then deriving its highest taxonomy rank using an in-house developed method, LastTaxa (Methods). If the majority of the MAGs in an unknown cluster are predicted to be in the same taxa at a certain rank, then the cluster can be assigned to that taxa. 9 of the 25 clusters are predicted to be Eukaryotes, and the remaining 16 clusters (56 MAGs) are all predicted to be Bacteria. Among these, 6 clusters were only classified at the superkingdom level, likely representing new phyla. 4 clusters were only classified at the phylum level, representing potential new classes. These clusters are listed in Table 1). The other clusters are better known, as they have close relatives in the reference database. They were missed because only subsets of reference genomes were used in the phylogenomics and clustering analyses.

**Table 1.** Top clusters representing candidate new taxa groups.

| cluster_label | rank | taxa | #MAGs | Note |
| --- | --- | --- | --- | --- |
| 598 | superkingdom | Bacteria | 4 | 35-40% proteins match Planctomycetes (phylum) |
| 730 | superkingdom | Bacteria | 6 | 12-20% proteins match Planctomycetes (phylum) |
| 758 | superkingdom | Bacteria | 4 | 36-93% proteins match Actinobacteria (phylum) |
| 1052 | superkingdom | Bacteria | 3 | 38-54% proteins match Actinobacteria (phylum) |
| 1059 | superkingdom | Bacteria | 3 |  |
| 1363 | superkingdom | Bacteria | 4 |  |
| 7 | phylum | Bacteroidetes | 3 | ~10% protein match Bacteroidetes bacterium (species) |
| 458 | phylum | Bacteroidetes | 3 |  |
| 816 | phylum | Chloroflexi | 3 | 44-66% proteins match Chloroflexia (class) |
| 1097 | phylum | Proteobacteria | 3 | one Actinobacteria |
| 1716 | class | Opitutae | 4 |  |

A complete list of the MAGs and their classification results can be found in Supplemental File 3. One striking observation from the Antarctic microbial communities is the presence of many large viruses. As our MAGs are at least 200kb, they likely represent giant viruses. When we construct a phylogenetic tree with 60 MAGs (28 from Bonney and 32 from Fryxell) with known giant virus references (Methods), they form several new branches, with 10 formed by 3 or more MAGs (Figure 3C). In addition, 6 MAGs co-cluster with these candidate giant viruses, raising the possibility that they may also be giant viruses. LastTaxa results provided independent support for three of them, one from Mimiviridae (BON.483) and two from Phycodnaviridae (FRY.48 and FRY.591), suggesting that Genome Constellation can provide a complementary approach to increase the sensitivity of giant virus detection.

## DISCUSSIONS

We presented a data-driven, genome-based framework to classify, cluster and visualize large numbers of microbial genomes based on their k-mer spectrum similarities. When applied to metagenomics, Genome Constellation can rapidly classify the taxa of medium-sized MAGs (200kb to 20Mb) if they have closely related references. *de novo* clustering the MAGs expands the ability of existing methods to do taxonomy classification, and facilities the identification of novel groups.

Whole-genome based taxonomy classification of the metagenome-assembled genomes (MAGs) is a computationally-intensive process as it often involves predicting all protein-coding genes, identifying phylogenetic markers and analyzing phylogenetic trees. For example, LastTaxa requires one to build databases using 500GB+ disk space, and it takes several minutes to predict the taxonomy of a single MAG. Although Genome Constellation is limited to only classifying genomes with closely related references, it can be used to quickly filter a large number of MAGs, and only pass on those without hits for further classification. This may dramatically reduce the computtation time required for analyzing large metagenome projects.

One extremely useful feature of Genome Constellation is its ability to cluster genomes, including those that lack known phylogenetic markers due to either phylogenetic divergence or assembly incompleteness. This was demonstrated in the analysis of MAGs predicted from the Antarctic metagenome dataset, as Genome Constellation identified several clusters of genomes that appear to originate from novel bacterial phyla.

There are a few limitations of Genome Constellation. First, its scalability in visualization is limited to networks with at most 100,000 nodes and 1 million edges. Scalable graph layout technologies such as Apache Spark-based Graph mapping (Jonker et al., 2017) could potentially be used for larger datasets. Second, although clustering and classification with pre-computed GSS is not limited by the size of the datasets, GSS computation with full spectra is still not very space- and time-efficient. A single full spectrum with a billion bits takes 128MB disk space, making it difficult to store the spectra of millions of genomes. This could be improved in the future by only storing a small set of representative references with minimum redundancy. Third, as the accuracy of taxonomy classification is dependent on the distribution of references in the GSS space, MAGs without any reference neighbor will not be classified. This problem will be alleviated by the rapid expansion of known references.

Genome Constellation can be used as a generic framework to form and visualize similar networks, such as genetic pathways, metabolites, etc. The framework only requires two prerequisites: a predefined scoring function or matrix so that the edge weight can be computed, and hierarchical annotation information for the nodes.

## ACKNOWLEDGMENTS

Zhong Wang, Dongwan Kang, Rob Egan, Shijie Yao, Jeff Froula, Volkan Sevim, Fredrik Schulz, and Tanja Wokye’s work was supported by the U.S. Department of Energy Joint Genome Institute, a DOE Office of Science User Facility, under Contract No. DE-AC02-05CH11231. Jackie Shay’s work was supported by JGI/UCM Genomics Summer Internship Program, Derek Macklin and Kayla McCue’s work was supported by DOE Computational Science Graduate Student Fellowhip, under contract No. DE-FG02-97ER25308, Rachel Orsini’s work was supported by DOE Mickey Leland Energy Fellowship. Rachael Morgan-Kiss, Wei Li and Chris Sedlacek were supported by a grant from NSF Office of Polar Programs (Award No. 1056396). DNA sequencing was funded by JGI Community Sequencing Project (CSP 1936) to Joan L. Slonczewski.

## AUTHOR CONTRIBUTIONS

DK, RE, and DM developed the GSS spectrum software. SY and ZW developed the web app. RMK, WL, CSW and JLS collected samples and isolated DNA. JF and VS performed the Antarctic metagenome analyses. HH and FS performed the phylogenetic tree analyses. JES designed and authored the Genome Constellation introduction video. KM, RO, DJB, SPB, CJS and ZW conducted additional data analyses. JLS and ZW designed and supervised the research. ZW, JLS and DM wrote the manuscript. All authors contributed to editing the manuscript.

